# Optimization of Genome-wide CRISPR Screens using dual-guide RNA Infection with Cas9 Electroporation (DICE)

**DOI:** 10.1101/2025.06.09.658313

**Authors:** Caitlin Purman, Charles Lu, Apexa Modi, Vinny Vijaykumar, Michael J. Flister, Anneke I. den Hollander, Sabah Kadri, Joshua D. Stender

## Abstract

Single-guide RNA lentiviral infection with Cas9 protein electroporation (SLICE) enables CRISPR screening in primary cell types that require transient Cas9 expression yet is limited by scalability and robustness. Here, we introduce dual-guide RNA infection with Cas9 electroporation (DICE), which expresses two guides from the same lentiviral construct that target the same gene. In genome-wide screens, DICE outperformed SLICE in defining essential genes and modulators of PD-L1 expression in IFN gamma activated THP1 cells. Collectively, these data demonstrate that DICE can be utilized for reduced-scale CRISPR screens in cell types with transient Cas9 protein expression without sacrificing screening quality.

## Background

CRISPR-Cas systems have emerged as a potent technology to provide readily programmable and highly efficient gene perturbations on single gene, multi-gene, and genome-wide scales. CRISPR-Cas dependent gene editing requires a guide RNA (gRNA) to recruit a Cas DNA nuclease (typically Cas9) to areas of homology in the genome where Cas9 creates a double-stranded break [1-3]. The imperfect repair of this lesion through non-homologous end joining results in an insertion or deletion (indel) of bases in the genome subsequently leading to a knockout of gene function [4]. While genome-wide CRISPR screens in immortalized cell lines have become routine, CRISPR screens in some biologically relevant models, like primary immune cells, present challenges that require modification of the standard approach [5-9]. Compared to immortalized cell lines, primary cells cannot be easily engineered to express Cas9 endogenously, and cell numbers are often limited, precluding the use of standard genome-wide CRISPR libraries which require hundreds of millions of cells for adequate gRNA coverage. To overcome the challenge of expressing Cas9 in primary immune cells, the single guide RNA (sgRNA) lentiviral infection with Cas9 protein electroporation (SLICE) method for CRISPR genome editing was recently developed [10]. SLICE enables CRISPR-mediated gene editing in cells that don’t express Cas9 protein and has been applied for genome-wide CRISPR screening in primary human T cells [10, 11]. While the development of the SLICE approach represents a major innovation for CRISPR screening in primary cells, to our knowledge, no head-to-head comparison of CRISPR screening with transient Cas9 (SLICE) versus constitutive Cas9 expression has been performed.

Here we compare CRISPR screening using the Brunello library [5] in THP1 cells with constitutively Cas9 expression or in THP1 cells requiring Cas9 protein electroporation and demonstrate that screening performance was more robust in THP1 cells constitutively expressing Cas9. To improve transient Cas9 screening approaches, we developed DICE, a multiplex gRNA lentiviral system expressing two gRNAs targeting the same gene from the same construct followed by Cas9 electroporation. We designed a miniaturized, genome-wide dual-Cas9 CRISPR library (21,034 constructs) and benchmarked it to the gold-standard Brunello library (76,441 constructs). We observe that, despite the reduction in library size, the dual-Cas9 library performs comparably to the Brunello genome-wide library in Cas9-expressing cells. Moreover, we show that the dual-Cas9 library (DICE) outperforms the Brunello library (SLICE) when Cas9 protein is introduced via electroporation. We thus present a novel approach and library that improves the outcome of genome-wide screens when Cas9 is exogenously introduced and drastically reduces the number of cells and resources needed to perform a genome-wide screen.

## Results

### Genome-wide SLICE has limited performance relative to traditional screening

Genome-wide CRISPR screens are routinely executed in immortalized cell lines, as these cells can be engineered to express Cas9 protein and expanded to essentially unlimited numbers; however, screening in cell lines does not always yield disease relevant targets. Therefore, there is an emphasis on developing approaches to enable CRISPR screening in primary cell models that are more likely to reflect disease biology and donor diversity. A hybrid CRISPR approach termed Single guide RNA Lentiviral Infection with Cas9 protein Electroporation (SLICE) was recently developed for pooled screening in primary human T cells wherein cells are transduced with a lentiviral CRISPR sgRNA library and Cas9 is subsequently introduced by electroporation [10]. Given that Cas9 activity within a cell persists for only 24-48 hours [12, 13] when Cas9 is introduced via electroporation and that most published screens allow 1+ weeks for genetic perturbation in Cas9-expressing cells, we sought to evaluate the global effect of Cas9 expression modes on CRISPR screen performance.

To evaluate the contribution of Cas9 expression on screening performance, we executed genome-wide screens for regulators of PD-L1 in THP-1 macrophages expressing Cas9 and in parental cells electroporated with Cas9 protein (**Figure 1A**) in response to IFN*α* stimulation. Briefly, we transduced Cas9-expressing or parental THP-1 cells with the Brunello genome-wide library at 500X coverage with a multiplicity of infection of 0.3 and selected for cells transduced with gRNAs. THP-1 parental cells expressing gRNAs were electroporated with Cas9 protein to induce genome editing following the SLICE protocol. Edited cells were then differentiated into macrophages with phorbol 12-myristate 13-acetate (PMA) and treated with IFN gamma to induce PD-L1 surface expression. Cells were FACS sorted for the top and bottom 10% of PD-L1 expressors. Thus, our screening workflow allowed us to assess library performance using both a viability and FACS-based readout.

**Figure 1:**
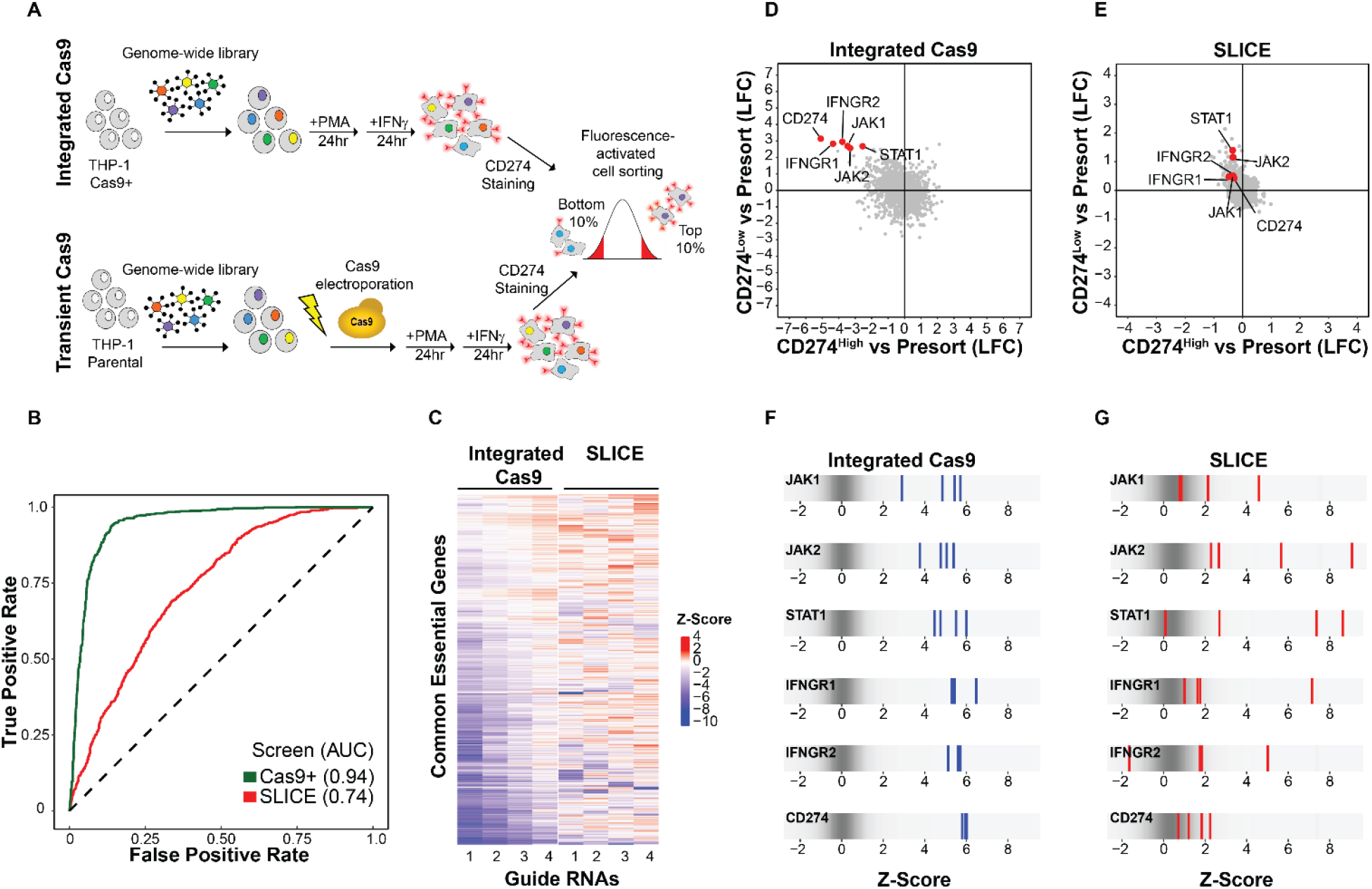
Performance of Brunello genome-wide CRISPR library with constitutive and transient Cas9 expression. A) Schematic overview of CRISPR screens to identify regulators of CD274 expression in THP-1 macrophages. Four screens were performed to enable a comparison of the dual-Cas9 and Brunello (single guide) platforms in the context of constitutive Cas9 expression and SLICE (electroporated Cas9). B) Single-gene receiver operating characteristic-area under the curve (ROC-AUC) plots derived from pan-essential and nonessential genes for Brunello screens in THP-1 or THP1-Cas9+ cells. C) Heatmap for the performance of each of four guide RNAs for common essential genes in THP1 cells with constitutive (integrated Cas9) or transient (SLICE) Cas9 expression. Data is displayed as Z-score normalized comparison between the guide counts in presort populations compared to pDNA levels. Comparison of gene enrichment in CD274^Low^ vs CD274^High^ sorted population in THP1 cells constitutively expressing Cas9 (D) or transient Cas9 expression (E). Density strip plots showing performance of each of the four guides targeting the core IFNγ signaling pathway in CD274^Low^ sorted population in THP1 cells constitutively expressing Cas9 (F) or transient Cas9 (G).

Receiver operating characteristic (ROC)-area under the curve (AUC) analysis of a logistic regression classifier that was trained with reference sets containing essential and nonessential genes was performed to estimate true positive and false positive rates for the Cas9 expressing cells and SLICE. This analysis revealed that the screen executed in Cas9 expressing cells (AUC=0.94) outperformed the screen with transient Cas9 expression (AUC=0.74) (**Figure 1B**). To further understand the difference in library performance between the stable and transient Cas9 screens, we compared the performance of the four gRNAs for each common essential gene in both screens. While the Z-score for guides targeting the same gene were well-correlated in the integrated Cas9 screen, this correlation was markedly reduced in the SLICE screen, indicating that the consistency of some individual guides is dependent on the mode and duration of Cas9 expression (**Figure 1C**).

FACS-based screening identified key regulators of IFNy-induced PD-L1 expression (**Figure 1D-E**) [14-16]. As expected, in Cas9-expressing cells, the core IFNy signaling pathway genes (JAK1/2, STAT1, IFNGR1/2) and CD274 itself were identified as top enriched genes in the PD-L1^low^ population (**Figure 1D**). In the SLICE screen, some of the positive controls were identified as significant hits including STAT1 (LFC=1.4) and JAK2 (LFC=1.2); however, strikingly, several core IFNy signaling pathway genes including IFNGR1 (LFC=0.479) IFNGR2 (LFC=0.504) and CD274 (LFC=0.429) were not significantly enriched in the PD-L1^low^ population (**Figure 1E**). The Z-scores of each of the four gRNA constructs for the known PD-L1 regulator genes were enriched to similar levels in the integrated Cas9 screen (**Figure 1F**). In contrast, in the SLICE screen, the variability in Z-scores for guides targeting the same gene increased dramatically, with some guides showing no enrichment (**Figure 1G**). Taken together, these results indicate gRNAs enrichment within genome-wide screens is highly dependent on the mode of Cas9 expression and delivery.

### Dual-guide RNA lentiviral infection followed by Cas9 electroporation (DICE) improves editing efficiency compared to SLICE

Multiplex Cas9 systems have emerged which express multiple sgRNAs which target the same gene or multiple genes from the same lentiviral construct. These approaches improve gene editing efficiencies compared to single guide systems [17]. Therefore, we reasoned that we could increase gene editing efficiencies in experimental contexts requiring nucleofection of Cas9 protein by leveraging a recently developed spCas9/dual gRNA approach wherein each lentiviral construct harbors two gRNAs targeting separate genes. This dual gRNA Cas9 (dual-Cas9) system features lentiviral constructs designed with separate U6-driven and H1-driven gRNA expression cassettes in opposing orientations. U6-VCR1 and H1-WCR3 promoter-tracr pairings were chosen based on superior efficacy and reduced recombination rate as previously reported [18]. Using this system, we designed lentiviral constructs with guides targeting both CD47 and CD63 expressed from both proximal and distal positions (**Figure 2A**). We transduced cells with both lentiviral constructs and then introduced Cas9 by electroporation. We observed greater than 75% knockout of both CD47 and CD63 with both construct orientations indicating that the U6 and H1 promoters express sgRNAs for similar gene editing efficiencies with transient Cas9 (**Figure 2A**).

**Figure 2:**
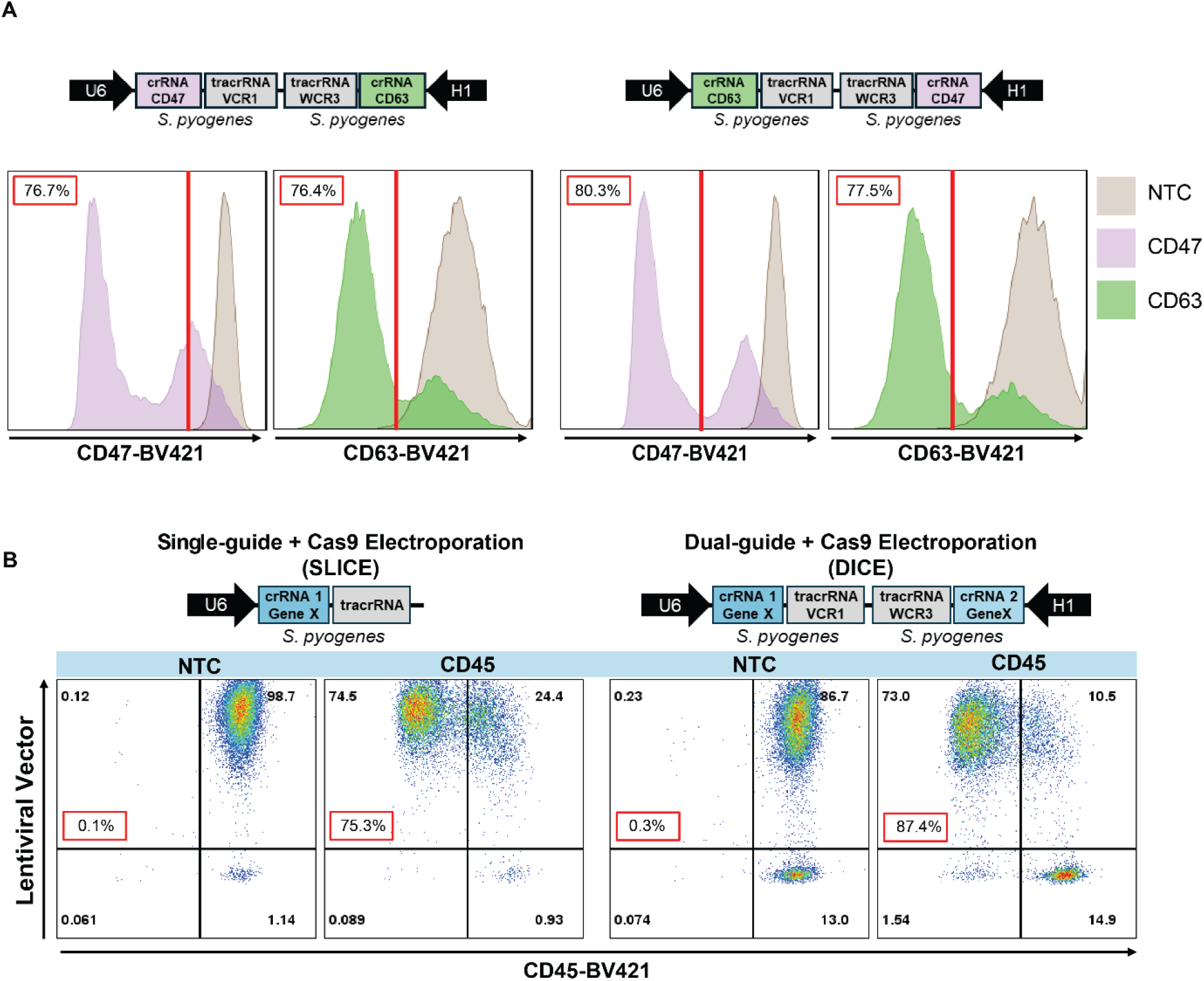
Development of DICE as strategy for multiplex gene editing. A) Schematic of dual-guide RNA expression cassettes targeting CD47 and CD63 (above). Evaluation of gene editing for CD47 and CD63 from guides driven by either the U6 and H1 promoters by flow cytometry (below). B) Comparison of targeting CD45 using single guides (SLICE, left) and dual-Cas9 constructs (DICE, right) using flow cytometry. CD45 levels were assessed by FACS at the indicated times post-transduction. Editing efficiency is indicated in the gray box in each FACS plot.

Previous reports have demonstrated that utilizing multiple guides targeting the same gene increases the probability of complete knockout by inducing deletions or indels [19, 20]. To increase the editing efficiency of the single-gene perturbation Cas9 system, we designed a dual-Cas9 construct containing two guide RNAs targeting the cell surface protein CD45 (**Figure 2B**). We assessed CD45 editing in cells transduced with lentiviral single- or dual sgCD45 constructs followed by electroporation of Cas9. Single-guide expression resulted in ∼75% CD45 editing efficiency. However, editing efficiency was augmented (∼87%) using the dual guide expression construct with transient Cas9. We thus present a dual-guide RNA lentiviral infection followed by Cas9 electroporation (DICE) approach which shows improved editing efficiency compared to SLICE.

### Design and benchmark of dual-Cas9 genome-wide library against the gold-standard Brunello library

To apply the DICE approach for genome-wide CRISPR screening, we designed a dual-Cas9 genome-wide CRISPR library (**Figure 3A**). Given the observed increase in editing efficiency with two guides per perturbation as opposed to one, we reasoned that we could efficiently target each gene with one dual-Cas9 construct as opposed to the four separate single-guide constructs per gene in the standard Brunello genome-wide library, reducing the total library size by 75%. Guide RNA sequences for the dual-Cas9 library were chosen based on the top two gRNAs for each target gene as predicted by Rule Set 3 [21], which incorporates variations in tracrRNA sequences and improves accuracy in predicting on-target efficiency over previous models. As the Brunello library was generated prior to the development of Rule Set 3, ∼70% of guides in the dual-Cas9 library are unique, while ∼30% are also found in the Brunello library **(Figure 3A)**.

**Figure 3:**
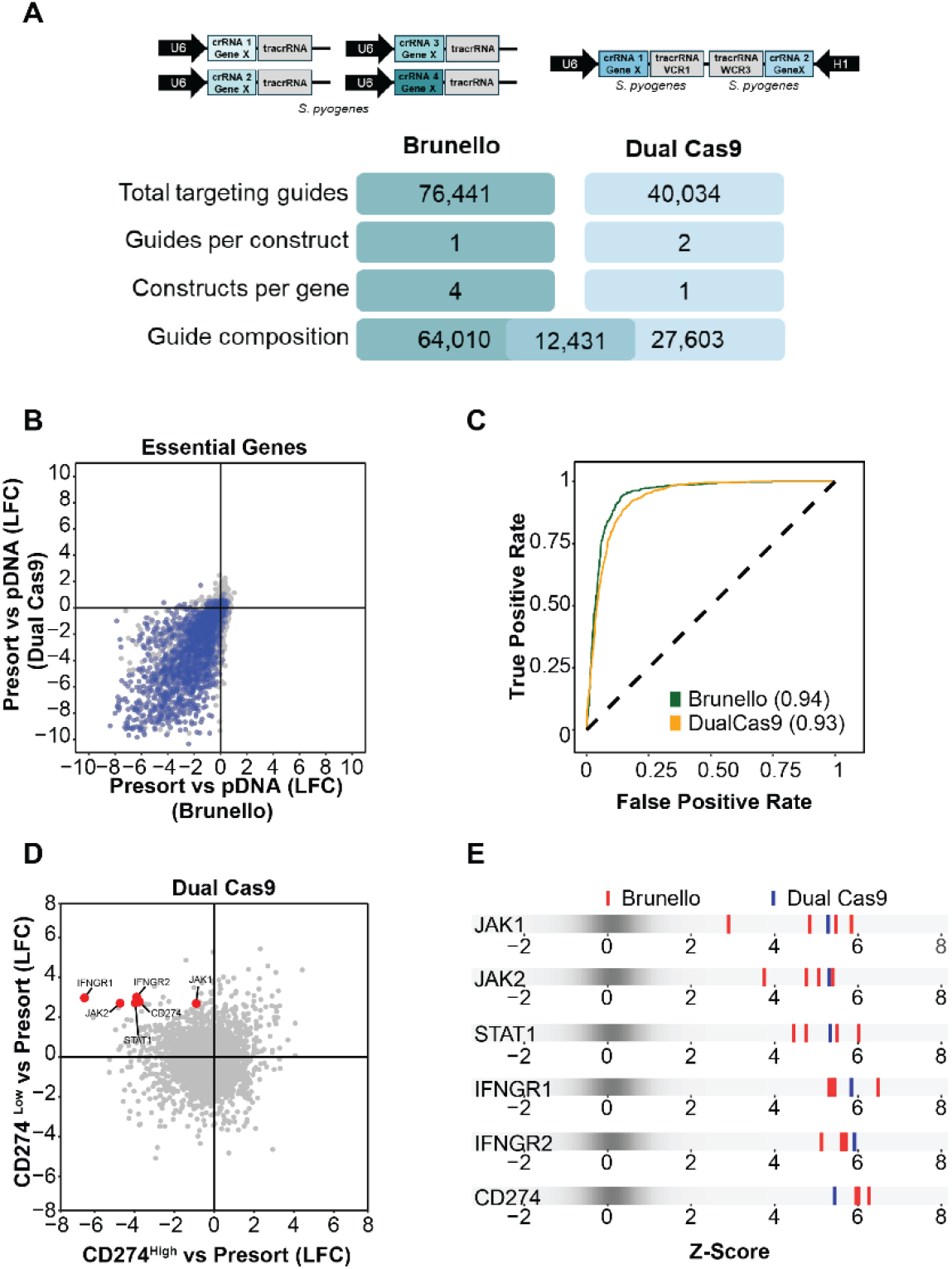
Miniature dual-Cas9 library provides comparable performance to Brunello library. A) Comparison and overlap of Brunello (single-guide) and dual-Cas9 genome-wide libraries. B) Scatter plot showing the correlation between lethality scores for essential genes in the Brunello and dual-Cas9 screens in THP-1 Cas9+ cells. C) Single-gene receiver operating characteristic-area under the curve (ROC-AUC) plots derived from pan-essential and nonessential genes for Brunello and dual-Cas9 screens in THP-1 Cas9+ cells. D) Scatter plots showing screening results for the dual-Cas9 screens. Each dot represents the median LFC of all sgRNAs targeting a particular gene. Dots for known CD274 regulator genes are labeled and shown in red. E) Density strip plots showing performance of guides targeting the core IFNγ signaling pathway in CD274^Low^ sorted population in THP1 cells constitutively expressing Cas9 and transduced with either the Brunello or Dual-Cas9 libraries.

### Dual-Cas9 and Brunello libraries perform comparably to identify common essential genes and PD-L1 regulators in Cas9-expressing THP-1 macrophages

We first compared the dual-Cas9 and Brunello library screening performance in THP-1 macrophages with stable Cas9 expression. The Pearson correlation for essential genes between the dual-Cas9 and Brunello was 0.806 and AUC score from ROC curve analyses indicated high and comparable performance between the two libraries in the detection of essential genes (Brunello, AUC 0.94; dual-Cas9 AUC 0.93) (**Figure 3B and 3C**). In addition to common essential genes, we assessed whether our dual-Cas9 library was able to identify known regulators of PD-L1 expression in response to IFNγ treatment (**Figure 1A and 3D)**. We plotted the median LFC of genes in our sorted PD-L1^low^ versus our PD-L1^high^ populations and found that all five of the known regulators as well as PD-L1 (CD274) itself were enriched in the PD-L1^low^ population and depleted in PD-L1^high^ population in dual-Cas9 screen (**Figure 3D**). When we compared the Z-score of each sgRNA construct in the Brunello (red) and dual-Cas9 (blue) libraries for five known regulator hits in our PD-L1^low^ population, we observed high enrichment and tight distribution of all constructs targeting each gene (**Figure 3E**). Collectively these results demonstrate that the novel dual-Cas9 library identifies regulators of PDL1 expression at similar robustness as the gold-standard Brunello library.

### DICE outperforms SLICE for genome-wide CRISPR screening

Given that our dual-Cas9 library performed comparably to the Brunello library in cells that stably express Cas9 and that gene editing efficiency is higher with the DICE approach compared to SLICE, we hypothesized that the dual-Cas9 library would outperform the Brunello library in screens where Cas9 expression is transient. The DICE platform would thus provide more robust gene editing to improve screening quality as well as enable genome-wide scale screens in cell models where scale is limited. We compared the performance of SLICE (Brunello) and DICE (dual-Cas9) genome-wide screens in our viability and FACS-based screens (**Figure 1A**). Depletion of essential genes was more robust in the DICE screen compared to SLICE, indicating improved editing efficiency with the dual-Cas9 library compared to Brunello (**Figure 4A**). Similarly, the DICE approach significantly outperformed the SLICE approach as shown by receiver operating characteristic (ROC)–area under the curve (AUC) analysis (**Figure 4B**). We next asked if our DICE screening platform was better able to identify known regulators of PD-L1 compared to SLICE. In the FACS-based screens, DICE identified all five known regulators of PD-L1 as significantly enriched in the PD-L1^low^ population whereas LFCs for known PD-L1 regulators were significantly lower in the SLICE screen, and PD-L1 itself was not identified as a statistically significant hit (**Figures 1E and 4C**). Moreover, the Z-score of each sgRNA construct in the Brunello (red) were generally lower than the guides in the dual-Cas9 (blue) library for known regulators of our PD-L1 (**Figure 4D**). As the sgRNA composition of the two libraries is not identical, performance of each library could depend on the repertoire of sgRNAs in each library and/or the dual compared to the single sgRNA library format. Of the five known regulators, both guides present in the dual-Cas9 library for JAK1 are also represented in the Brunello library. Interestingly, while all three constructs performed comparably in the integrated Cas9 screens, neither individual guide constructs in the Brunello library performed as well as the dual guide construct in the transient Cas9 screens, pointing to a likely increase in editing efficiency with two sgRNAs verses one when Cas9 is not stably expressed (**Figure 3E** and **Figure 4D**). These data suggest that the DICE approach significantly outperforms the SLICE in viability- and FACS-based CRISPR screens.

**Figure 4.**
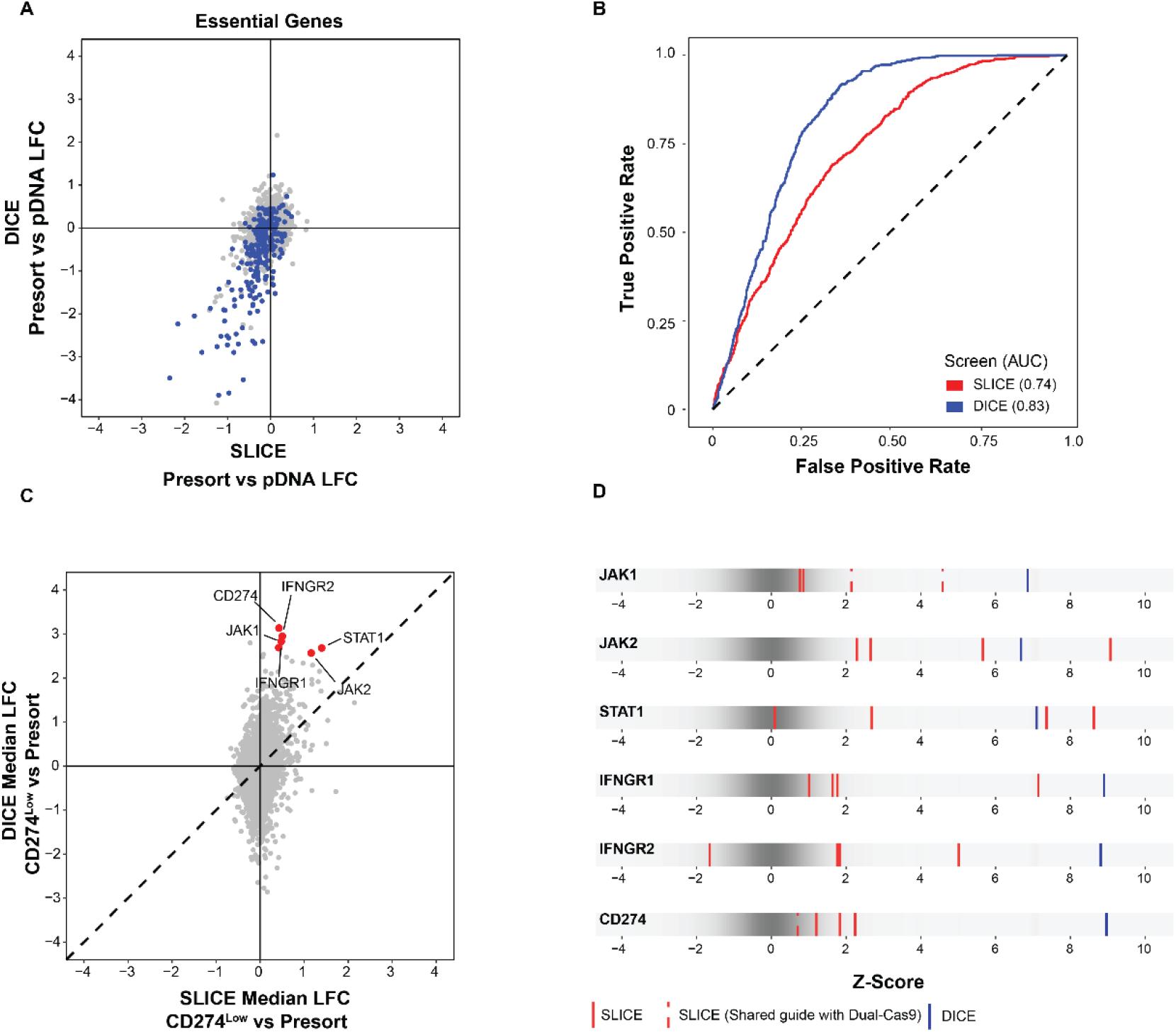
dual-Cas9 outperforms Brunello to identify essential genes and PD-L1 regulators in THP-1 macrophages when Cas9 is introduced transiently. A) Scatter plot showing the correlation between lethality scores for common essential genes in SLICE vs DICE screening formats where the two Brunello library guides overlap with Dual-Cas9 library. B) Single-gene receiver operating characteristic-area under the curve plots derived from pan-essential and nonessential genes for Brunello and dual-Cas9 in Cas9+ THP1 cells or in either the SLICE or DICE screening format. C) Comparison of SLICE vs DICE for CD274^low^ populations. The core IFN pathway genes are highlighted in red. D) Density strip plots showing performance of guides targeting the core IFNγ signaling pathway in CD274^Low^ sorted population in SLICE (Red) and DICE (Blue). Guides that are shared between Brunello and Dual-Cas9 are dotted red lines.

## Discussion

In this study, we present a new approach for improved CRISPR editing using single-target, dual sgRNA expressing lentiviral constructs and Cas9 protein electroporation. We designed a genome-wide dual sgRNA (dual-Cas9) library wherein each gene is targeted by two separate sgRNAs expressed from a single construct, reducing the library size by 75% compared to the gold standard Brunello genome-wide library. We compared the performance of the dual-Cas9 genome-wide CRISPR library to the Brunello library using both viability- and FACS-based screening readouts. We confirm that the dual-Cas9 library performs comparably to the Brunello library in cells expressing Cas9 but significantly outperforms the Brunello library in screens where Cas9 is delivered exogenously. The observation that both libraries perform similarly in the context of constitutive Cas9 suggests that genome-wide screens can be scaled down by 75% without missing valuable biological information. Miniature libraries can enable genome-wide screening in cell types where cell numbers are limited such as primary immune and neuronal cells. At the same time, the dual-Cas9 library, which expresses two sgRNAs targeting the same gene, provides a performance advantage over Brunello library in screens where Cas9 must be introduced by electroporation. A potential explanation for improved screening quality with the dual-Cas9 despite reduced library size is that under transient expression, Cas9 protein has twice as many opportunities to edit the genome in each cell with the dual-Cas9 system compared to a single guide library. This explanation is supported by the tight distribution of LFCs for Brunello and Dual-Cas9 library constructs targeting known PD-L1 regulators in screens in THP-1 Cas9+ cells, but relative loss of enrichment of many Brunello constructs compared to Dual-Cas9 constructs in the SLICE screens. In SLICE experiments key members of the JAK/STAT pathway were not identified as significant hits when using the Brunello library suggesting that single guide libraries have the potential to miss important regulators of the targeted biology. Our dual-Cas9 platform fills critical gaps in CRISPR screening capabilities to enable improved screen performance when Cas9 is transient and reduces genome-wide screening scale, both of which are critical for screening in primary human immune and neuronal cells.

## Conclusions

This study presents a novel dual-Cas9 genome-wide CRISPR library consisting of one gRNA construct per gene and compares its performance to the four-construct-per-gene Brunello library. In both viability- and FACS-based screens, the dual-Cas9 library performs similarly to Brunello when Cas9 is constitutively expressed and outperforms Brunello when Cas9 is delivered exogenously. The dual-Cas9 platform enables screening with 75% fewer resources without compromising quality, making it ideal for primary cells with limited numbers. Moreover, its superior performance in contexts with exogenous Cas9 highlights the potential of the dual-Cas9 library to capture biologically valuable information missed by single-guide libraries.

## Methods

### Cell culture

THP-1 cells were maintained in RPMI 1640 (Gibco) supplemented with 10% heat inactivated fetal bovine serum (FBS, Gibco), 1% Pen/Strep (Gibco), 1% L-glutamine (ThermoFisher Scientific, Cat# 25030081), 1% Sodium pyruvate (ThermoFisher Scientific, Cat# 11360070), 1% HEPES (ThermoFisher Scientific, Cat# 15630-080), and 0.05 mM 2-mercaptoethanol (ThermoFisher Scientific, Cat# 21985023) in a 37°C humidified 5% CO_2_ incubator. Cells were differentiated by the addition of 10 ng/ml phorbol 12-myristate 13-acetate (Sigma, P1585) for 24 hours and treated with 10 ng/ml IFN gamma (PeproTech, Cat# 300-02) for 48 hours.

### Generation and characterization of Cas9-stable cells

Cas9-stable cells were generated by infecting parental cell lines with a lentiviral construct expressing Cas9 and blasticidin-resistance gene in 12-well plates at 1000 x g for 1 h, in the presence of 8 μg/μL polybrene (Sigma-Aldrich). Plates were then returned to 37 °C with 5% CO_2_. Cells were incubated overnight and then selected by 10 μg/ml blasticidin (Thermo Fisher Scientific). The activity of Cas9 was confirmed to be greater than 75% through introducing guides targeting CD274, CD45, and confirming loss of cell surface expression compared to a non-targeting control by flow cytometry.

### Design of dual-Cas9 library

Guide RNAs for each human genes were designed using Broad’s Rule Set 3 [21] and are listed in Supplementary Table 1. The oligo library was synthesized at Twist Biosciences and cloned using a two-step process where the guides were cloned into pvAA0017 backbone followed by introduction of the TRACR RNA cassettes. The TRACR RNA sequences downstream of the U6 and H1 promoters are VCR1 and WCR3 respectively [18].

### Genome-wide CRISPR screening

For the Brunello library transductions, 120 million THP1 cells or THP1 stably express Cas9 were transfected with the sgRNA lentiviral library at multiplicity of infection (MOI) = 0.3 in three biological replicates. The Brunello CRISPR-Cas9 knockout library contains 76,441 sgRNAs targeting 19,114 genes with 1,000 non-targeting controls. For Dual-Cas9 library transductions, 30 million THP1 cells or THP1 stably express Cas9 were transfected with a lentiviral library at multiplicity of infection (MOI) = 0.3 in three biological replicates. Following transduction, transduced cells were selected with puromycin (2 μg/ml, ThermoFisher Scientific, Cat# A1113802) and dead cells were removed (Miltenyi Biotec, Cat# 130-090-101). For SLICE experiments, 800 pmol Cas9 was nucleofected per 20E6 cells using the Expanded T cell program 2 on MaxCyte STx nucleofector. Cells were then differentiated with PMA and treated with 10 ng/ml IFN gamma for 24 hours. 10 (dual Cas9) and 40 million (Brunello) cells were collected as presort for genomic DNA extraction for each replicate. Greater than 40 or 10 million cells per replicate for Brunello or Dual-Cas9 experiments respectively were then processed and stained for anti-Human PDL1 cell surface expression (Biolegend, Cat# A1113802) at 10 million cells/ml according to manufacturer’s recommendation. Cell populations expressing the top and bottom 10% PDL1 levels were enriched through FACS sorting using Aria Fusion (BD Biosciences). At least one million PDL1^low^ and PDL1^high^ cells were collected for genomic DNA extraction using Quick-DNA FFPE Kit (Zymo Research). PCR reactions containing up to 10 μg genomic DNA in each reaction were performed using Titanium Taq DNA Polymerase with primers to amplify the sgRNAs. Samples were then purified with SPRIselect beads, mixed and sequenced by 75-bp single-end sequencing for Brunello libraries and 2×150 bp paired-end sequencing for dual-Cas9 libraries.

### CRISPR screen analysis

Pair-end sequencing data were demultiplexed and paired reads each with a sgRNA were quantified using custom Perl scripts. Given that the position of the sgRNA could vary per read, the primer sequences flanking the sgRNA were considered. Sequences in between the flanking primers were extracted and then compared to sequences in the sgRNA library. Only sequences with no mismatches were used in the calculation of guide-level read counts. Due to possibilities of recombination events, only pair of sgRNAs that maps to the same gene or only 1 sgRNA were identified in each read-pair were used for downstream analysis. Overall, samples with a minimum of 75% reads mapping to a sgRNAs were considered for further analysis. Through the *edgeR* Bioconductor package, reads were normalized between samples using the trimmed mean of M values (TMM) method [22]. Differential analysis of guide level counts was performed using the *limma* R Bioconductor package [23]. Gene level effects were considered as the median log fold change of the four guides associated with each gene. Statistically significant genes were identified by aggregating differential sgRNAs ranked consistently higher at gene level using robust rank aggregation (RRA) method. Benchmarking the performance of the various CRISPR platform was described in [24].

## Acknowledgements

We thank David Root, Olivia Bare, John Doench, and other members of the Genetic Perturbation Platform of The Broad Institute (Cambridge, MA) for the Brunello library as well as the design and cloning of the dual-Cas9 sgRNA library. We thank Jing Wang, Si Wu and Fedik Rahimov for critical review of the manuscript.

## Author contributions

C.P. C.L., M.J.F, and J.D.S conceived, designed and led the study and drafted the manuscript. C.P., V.V., performed experiments, C.L., A.M., C.P., J.D.S., analyzed the data, S.K., M.J.F., A.I.D, provided resources and critical feedback. All authors reviewed the manuscript, provided feedback, and approved the final version of the manuscript.

## Declarations

All authors are or were employees of AbbVie at the time of the study. The design, study conduct, and financial support for this research were provided by AbbVie. AbbVie participated in the interpretation of data, review, and approval of the publication. No honoraria or payments were made for authorship.

